# A novel pyridoindole improves the recovery of residual hearing following cochlear implantation after a single preoperative application

**DOI:** 10.1101/2024.02.14.580226

**Authors:** Michael Nieratschker, Erdem Yildiz, Matthias Gerlitz, Sujoy Bera, Anselm J. Gadenstaetter, Anne-Margarethe Kramer, Monika Kwiatkowska, Pavel Mistrik, Lukas D. Landegger, Susanne Braun, Reimar Schlingensiepen, Clemens Honeder, Christoph Arnoldner, Hans Rommelspacher

**Affiliations:** Department of Otorhinolaryngology, Head and Neck Surgery, Medical University of Vienna, Vienna, Austria; AudioCure Pharma GmbH, Berlin, Germany; Center for Biomedical Research and Translational Surgery, Medical University of Vienna, Vienna, Austria; MED-EL Medical Electronics, Austria

**Keywords:** Hair cell preservation, Inner ear, Hearing preservation, Cochlear implantation, Synaptopathy

## Abstract

Sensorineural hearing loss (SNHL) is the most common sensory deficit worldwide. Due to the heterogeneity of causes for SNHL, effective treatment options remain scarce, creating an unmet need for novel drugs in the field of otology. Cochlear implantation (CI) currently is the only established method to restore hearing function in profound SNHL and deaf patients. The cochlear implant bypasses the non-functioning sensory hair cells (HCs) and electrically stimulates the neurons of the cochlear nerve. CI also benefits patients with residual hearing by combined electrical and auditory stimulation. However, the insertion of an electrode array into the cochlea induces an inflammatory response, characterized by the expression of pro-inflammatory cytokines, upregulation of reactive oxygen species, and apoptosis and necrosis of HCs, putting residual hearing at risk.

Here, we characterize the effects of the small molecule AC102, a pyridoindole, for its protective effects on residual hearing in CI. We show that AC102 significantly preserves hearing thresholds across the whole cochlea and confines the cochlear trauma to the directly mechanically injured area. In addition, AC102 significantly preserves auditory nerve fibers and inner HC synapses throughout the whole cochlea. AC102s effects are likely elicited during the inflammatory phase of electrode insertion trauma (EIT) and mediated by anti-apoptotic and anti-inflammatory properties, as uncovered by an *in vitro* assay of ethanol induced apoptosis and evaluation of mRNA expression of pro-inflammatory cytokines in an organotypic *ex vivo* model of EIT.

The results in this study highlight AC102 as a promising compound for the attenuation of EIT during CI. Moreover, as the inflammatory response in cochlear implantation shares similarities to other etiologies of SNHL, a beneficial effect of AC102 can be inferred for other inner ear conditions as well.

## Introduction

Hearing loss is the most common form of sensory deficit worldwide, with estimates by the World Health Organization predicting that over 450 million people will suffer from disabling hearing loss by 2030 [1]. Sensorineural hearing loss (SNHL) results from damage to the cochlear sensory hair cells (HC) and their associated neural connections, caused by a variety of factors such as aging, noise exposure or ototoxic drugs [2]. SNHL is irreversible due to a lack of regenerative capacity of the cochlear HCs, and given the heterogeneity of etiologies, effective treatment options remain scarce. Apart from the recently approved sodium-thiosulfate for cisplatin-induced ototoxicity [3,4], there is currently no FDA approval for any etiology of SNHL, highlighting the need for novel drugs in the field of otology.

AC102 is a small lipophilic pyridoindole currently under investigation for treating sudden SNHL [5]. In an animal model of noise trauma, AC102 significantly preserved auditory thresholds by protection of HCs and inner HC synapses, likely mediated by a reduction of reactive oxygen species (ROS), temporary upregulation of ATP, and stimulation of neurite outgrowth [6]. Further, a structurally related compound to AC102 named 9-methylpyridoindole (9MP) was found to exert anti-apoptotic and anti-inflammatory effects by regulation of pro-inflammatory cytokines and reduced buildup of ROS [7–9]. AC102s structural difference to 9MP further enhances its efficacy and increased its half-life in the perilymph of guinea pigs 1.75-fold [10,11]. Owing to these properties, AC102 is emerging as a promising treatment for inner ear disorders.

In patients with profound SNHL or deafness, cochlear implantation (CI) is currently the only established method to restore hearing function. The cochlear implant is a neural prosthesis that is surgically inserted into the snail-shaped cochlea of the inner ear. Through electrical stimulation, it bypasses the non-functioning auditory HCs and directly stimulates the spiral ganglion neurons (SGNs) of the cochlear nerve. Sound transduction in the cochlea is tonotopically organized, where each frequency is mapped to a specific position, ranging from high frequencies at the base to low frequencies at the apex. In patients with residual hearing in the low frequencies, technical and surgical advancements allow a combined electrical and acoustic stimulation (EAS) [12,13]. Here, the cochlear implant provides electrical stimuli in the damaged basal part of the cochlea, while the preserved apical area is stimulated with amplified sound, resulting in a synergistic improvement of speech discrimination, musical appreciation, speech-in-noise perception, and overall quality of life [14–17]. However, the loss of residual hearing in EAS-CI recipients is common [14,17–19], and is primarily caused by an immediate trauma during CI, followed by a subsequent inflammatory response [20–22]. Even a cautious insertion with a flexible electrode array can elicit an immune response, which drives the expression of pro-inflammatory cytokines and initiates the buildup of ROS [23–25]. This ultimately leads to deterioration of cochlear cells through activation of pro-apoptotic pathways, necrosis, and necrosis-like cell death [26,27]. A wound healing response then induces deposition of fibrocytes and cochlear fibrosis [24,27], further hindering the wave propagation towards the cochlear apex [28].

Pharmacological treatment of the inflammatory cascade during electrode insertion trauma (EIT) has been of great interest in recent years [29]. Due to their potent anti-inflammatory properties, glucocorticoids (GCs) could successfully preserve residual hearing and cochlear structures following CI as demonstrated in animal studies [30–32]. However, translation into human trials observed only minor protective effects [31,33]. Similarly, other pharmaceutical compounds acting on specific targets along the inflammatory pathway, including specific inhibitors of apoptosis, anti-inflammatory compounds, and antioxidants failed to find favorable outcomes to reduce EIT so far [34–36].

In the present study, we evaluated the effects of a single preoperatively and locally applied AC102 hydrogel for the preservation of residual hearing in EAS-CI. AC102s efficacy was investigated by means of electrophysiological measurements as well as histological analysis in an animal model of CI. We further elucidated its mode of action in an *in vitro* model of apoptosis and in an *ex vivo* model of EIT. Our results highlight AC102 as a promising small molecule for the attenuation of EIT and support further investigation as a causal treatment option in SNHL.

## Materials and methods

### 1. Animal housing

All animal experiments were approved by the local animal welfare committee and the Austrian Federal Ministry for Science, Research and Economy (66.009/0094-V/3b/2019). In total, 18 female Mongolian gerbils (Charles River Laboratories, Sulzfeld, Germany) aged between 30 and 60 days were used.

### 2. Gel administration and cochlear implantation

*Experimental design*: Unilateral CI was performed in 18 female Mongolian gerbils. 24 hours prior to implantation, an AC102 hydrogel (AC102) or unloaded vehicle hydrogel (Vehicle) was surgically applied in the round window niche. Auditory brainstem responses (ABR) were used to objectively measure auditory function approximately one week before hydrogel administration to ensure normal hearing prior to inclusion in the experiments. Auditory compound action potentials (CAP) were measured before hydrogel injection, following CI, and on postoperative days 3, 7, 14, and 28 (**Fig. 1**). Impedance measurements of the electrode were performed on the same days following CI. At the study endpoint, temporal bones were extracted, cochlear whole mounts were prepared and immunofluorescence staining using various antibodies was performed. Inner ear HCs, ANFs, presynaptic ribbons and postsynaptic ANF terminals of the IHCs were evaluated over the whole cochlear length.

**Fig. 1.**
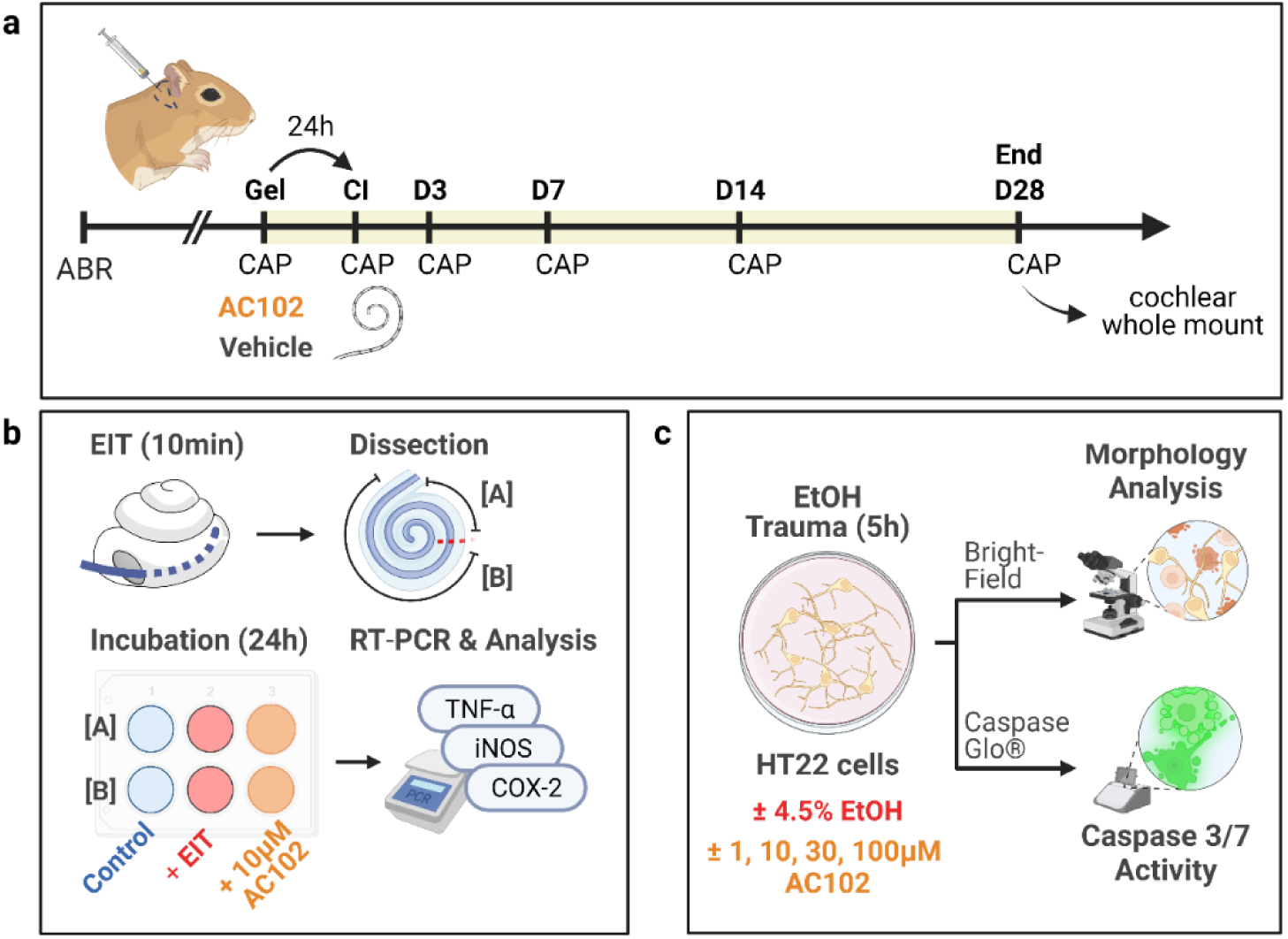
Schematic representation of the experiments. **a.** The effect of a locally applied AC102 hydrogel 24 hours prior to cochlear implantation (CI) was compared to a Vehicle-treated control group. Normal auditory function was assessed one week prior to inclusion in the study by auditory brainstem responses (ABR). 24 hours prior to cochlear implantation (Gel), a baseline compound action potential (CAP) measurement was carried out before the administration of 10 μL of the respective hydrogel via a retroauricular approach. On day 0 (CI), cochlear implantation was performed, and postoperative CAP and impedance measurements were carried out. Auditory function was assessed on days 0, 3, 7, 14 and 28 by CAP and impedance measurements. After the last experimental timepoint (End), animals were sacrificed, and their cochleae extracted for whole mount analysis. **b**. Schematic representation of an *ex vivo* EIT model. Cochleae of P4-C57BL/6 mice were extracted, and trauma induced by inserting a 2-0 suture into the round window and incubated for 10 minutes (EIT). Cochleae were then dissected into a basal [A] and middle to apical [B] part, and incubated for 24 hours in culture medium with or without 10 µM of added AC102. mRNA expression of inflammatory enzymes was then evaluated by RT-qPCR. **c.** Schematic representation of cell damage and apoptosis assays. HT22 cells were damaged with 4.5% ethanol (EtOH) for 5 hours and treated with varying concentrations of AC102. Cell damage was quantified by changes in cell morphology (round cells), and apoptosis was evaluated by Caspase 3/7 activity with a commercial kit as well as immunohistochemistry. Created with BioRender.com

*Anesthesia and supportive care:* Hydrogel application and CI surgery were both performed under general anesthesia using subcutaneously (s.c.) applied medetomidine (0.3 mg/kg BW), midazolam (1 mg/kg BW), fentanyl (0.03 mg/kg BW), and ketamine (10 mg/kg BW). To maintain anesthesia during surgery, 33% of the initial anesthetic dose was resupplied every 30 minutes. Lidocaine (2 mg/kg BW) was injected s.c. in the surgical area prior to skin incision. Perioperatively, 7 mg/kg BW Enrofloxacin was injected s.c. once daily for three days starting on the day of hydrogel application. For auditory testing under sedation, the same drug regimen without fentanyl was used. After the experiments, anesthesia was partially antagonized using atipamezole (1 mg/kg BW) and flumazenil (0.1 mg/kg BW).

*Hydrogel formulation and administration:* Both, AC102 (AudioCure Pharma, Berlin, Germany) and Vehicle hydrogel were provided in a readily injectable thermosensitive formulation, containing 12 mg/ml AC102. The formulations were stored at 4°C until use and resuspended by vortexing prior to injection. The hydrogel was applied via a retroauricular approach [37]. The postauricular region of the right ear was shaved and subsequently disinfected with povidone-iodine solution. Three millimeters behind the pinna, a two cm long skin incision was cut, the postauricular muscles were bluntly dissected, and the bony wall of the bulla exposed. The bulla was carefully opened, and the round-window (RW) niche identified. A teflon-insulated gold wire (Goodfellow, Bad Nauheim, Germany), which served as the CAP recording electrode, was hooked to the bony ridge of the RW niche and temporarily fixed with Histoacryl® glue (Braun Melsungen, Melsungen, Germany). Afterwards, a baseline CAP measurement was carried out. Following auditory measurement and removal of the gold wire, 10 µL of the compound (0.12 mg AC102 or Vehicle) were applied to the RW niche using a YOU-1 micromanipulator (Narishige, Tokyo, Japan), and Hamilton syringes (Hamilton, Bonaduz, Switzerland) with blunt 29G needles. To ensure an appropriate solidification of the hydrogel, a tilted animal position was maintained for another ten minutes during wound closure. The bulla was then temporarily closed by placing a silastic foil (Invotec International Inc., FL, USA) over the bony opening and the edges sealed with Histoacryl® glue. The wound was sutured in a two-layered fashion using absorbable sutures (Vicryl® 4-0, Ethicon, Somerville, NJ, USA).

*Cochlear implantation:* 24 hours following hydrogel application, CI was performed. Under general anesthesia, two 4mm titanium bone-screws (#19010-10, FST, Heidelberg, Germany) were superficially screwed into the skull bone as an anchor to fix the implant connector on the animal’s skull. The retroauricular wound was reopened, and residual hydrogel was thoroughly removed from the bulla by rinsing with sterile sodium chloride under careful suction. The RW niche was widened using a 0.5-mm tapered diamond burr, and a gold wire recording electrode was again hooked at the RW overhang. A custom-designed, 4mm long CI electrode (MED-EL, Innsbruck, Austria) with two electrode contacts (basal and apical) was then inserted twice via the RW in order to elicit the desired amount of EIT. The maximum insertion depth tonotopically corresponds to 5.12 kHz [38]. The electrode connector was fixed onto the animals’ vertex, and the CAP wire was soldered to the connector. The bulla was closed permanently using Paladur® dental cement (Kulzer GmbH, Hanau, Germany), the wound was sutured in a two-layered fashion, and the connector together with the screws was securely fixed to the skull with dental cement.

### 3. Electrophysiology

*Auditory brainstem response and compound action potential:* The general setup for auditory testing is described in detail in a previous publication [39]. In brief, audiometric measurements were performed in a sound-proof chamber (mac-2; Industrial Acoustics Company, Niederkrüchten, Germany) with a DT-48 loudspeaker positioned 3cm from the tested ear and a K2 microphone (Sennheiser, Wedemark, Germany) placed directly above the pinna for calibration. The contralateral ear was occluded with a wax earplug. Auditory potentials were measured via a custom-made setup, including a PC system equipped with a multifunction I/O card and AudiologyLab software (Otoconsult, Frankfurt am Main, Germany). ABR and CAP thresholds were recorded in 5 dB steps in a frequency range of 0.5-32 kHz following the stimulation of clicks and tone-bursts (3ms duration, 1ms rise/fall), at 1- or 3-steps per octave for ABRs and CAPs, respectively. Threshold shifts were calculated for every frequency and timepoint by subtracting the baseline threshold from the postoperative result. CAP thresholds were determined visually for all treatment groups in a blinded manner. Auditory thresholds were defined as the sound pressure level (dB SPL), for which the CAP wave-1 could be clearly identified and which persisted or increased at higher stimulation intensities. CAP threshold shifts were grouped into four frequency ranges to represent the insertion trauma-affected cochlear regions where the electrode was actually located (6.3-12.6 kHz and 16-32 kHz) and the regions apical of the electrode insertion depth (0.5-1.5 kHz and 2-5 kHz, see **Fig. 2**).

**Fig. 2.**
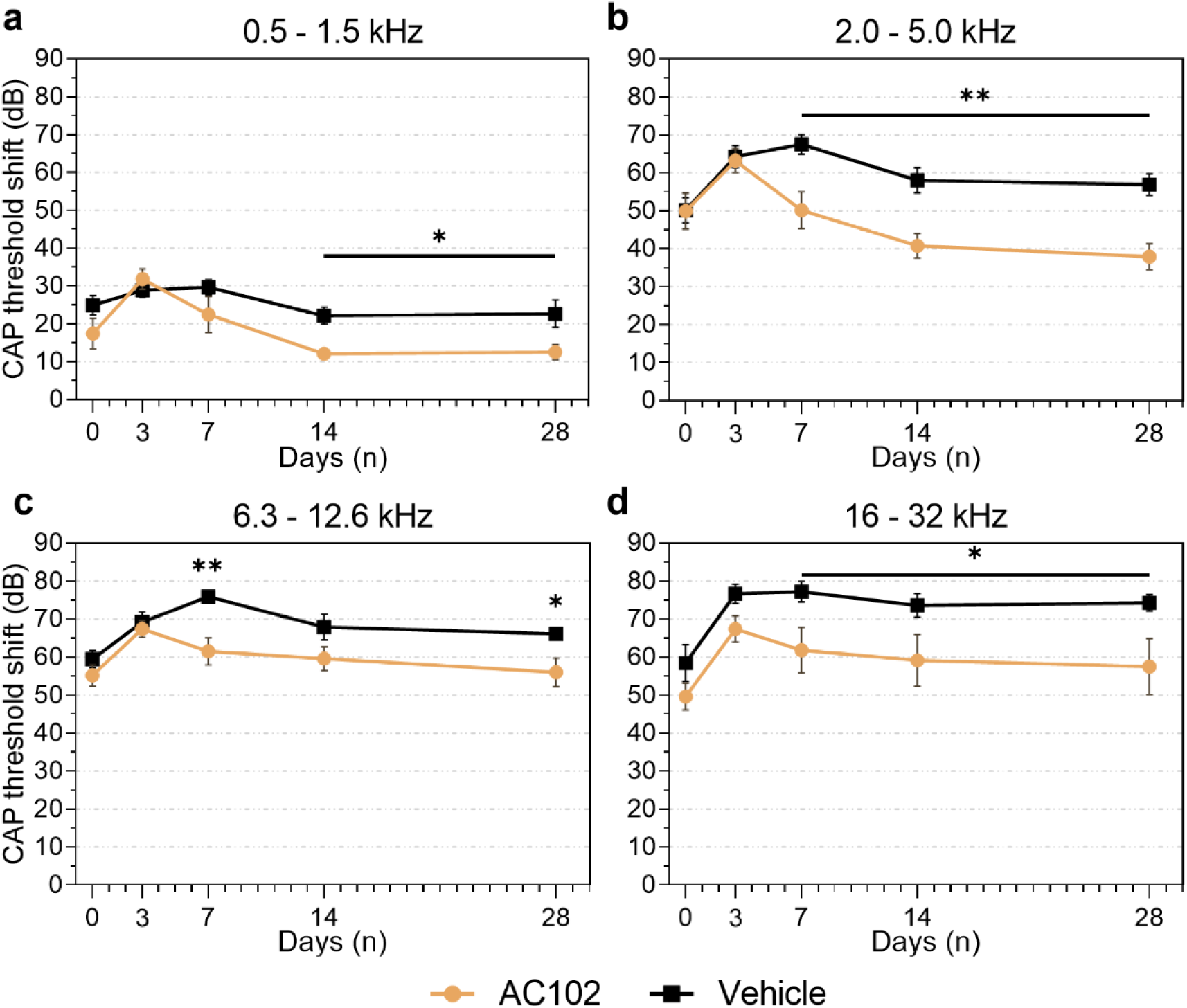
Compound action potential (CAP) threshold shifts (dB) over the course of 28 days after cochlear implantation, treated with AC102 (n = 9, orange) or Vehicle hydrogel (n = 9, black) 24 hours prior to implantation. Measured frequencies were grouped into (a) 0.5-1.5 kHz, (**b**) 2.0-5.0 kHz, (**c**) 6.3-12.6 kHz, and (**d**) 16-32 kHz. Error bars = SEM, * = p < 0.05, ** = p < 0.01

*Impedance measurements:* Impedance measurements were carried out on the same time points as auditory testing as a marker for intracochlear tissue growth. Impedances were evaluated via a MAX-Box and Pulsar I100 service implant in Maestro software version 5 (MED-EL). Beginning on day 3, measurements were performed after 15 min of electrical stimulation (pulse width: 35.42 ms, pulse rate: 750/s, engraved power: 452.1 mA) to simulate conditioning as is used in clinical practice. Impedances of both the apical and basal electrodes were repeatedly measured 5 times and averaged.

### 4. Cochlear whole-mount analysis

At the last experimental timepoint, animals were transcardially perfused with buffered 4% paraformaldehyde (PFA; Sigma Aldrich, Seelze, Germany), and their inner ears were extracted. Under microscopic vision, the cochlear implant and stapes were carefully removed, and a small hole was drilled into the cochlear apex in order to create basal and apical openings. Inner ears were flushed and subsequently immersed in PFA for 24 hours followed by decalcification in 4% ethylenediaminetetraacetic acid (EDTA) for 4 days. Cochleae were then dissected as whole mounts into 4-6 pieces in total. Care was taken to preserve the whole cochlear length for later frequency map determination.

For antibody staining, the specimens were immersed in blocking solution containing 1% Triton-X and 5% normal goat serum (NGS) in phosphate-buffered saline (PBS) for 1 hour. Tissue was then incubated in primary antibody solution overnight at 37°C. Rabbit-IgG anti Myosin-7A @ 1:200 (Enzo Life Sciences, #ALX-210-227-R200) was used to label HCs, Chicken-IgY anti NF200kD @ 1:250 (Millipore, #AB55339) was used to label ANFs, Mouse-IgG1 anti CtBP2 @ 1:200 (BD Transduction Labs, #612044) and Mouse-IgG2a anti GluA2 @ 1:2000 (Millipore, #MAB397) were used to label presynaptic IHC ribbons and postsynaptic terminals, respectively. Following three washing steps in PBS, specimens were transferred into secondary antibody solution for one hour at room temperature. Alexa Fluor (AF)-conjugated antibodies matched to the above-mentioned primaries were chosen at four different wavelengths at concentrations of 1:500 (Goat anti Chicken-IgY AF-647 and Goat anti Rabbit-IgG AF-405) or 1:1000 (Goat anti Mouse-IgG1 AF-568 and Goat anti Mouse-IgG2a AF-488). After three more washing steps in PBS, specimens were embedded in Prolong Gold Antifade (Life Technologies, #P36930) and stored at room temperature for 24 hours until imaging.

Confocal imaging was performed using a Nikon Ti Eclipse confocal microscope (Nikon, Tokyo, Japan). Cochleae were imaged at 20x magnification and each cochlear turn was stitched using the ImageJ Plugin Mosaic-J [40]. Using the ImageJ Plugin “Measure Line”, outer HCs (OHC) and inner HCs (IHC) were grouped into 20 equal-sized divisions (https://www.masseyeandear.org/research/otolaryngology/eaton-peabody-laboratories/histology-core). IHCs and OHCs were evaluated per section along the whole cochlear length and expressed as mean percent compared to the contralateral side (= 100%). Furthermore, a cochlear frequency map was calculated for each specimen [38]. IHC synapses and type-II ANFs were evaluated in one-octave steps between 0.5 and 32 kHz. Type-II ANFs were investigated spanning 100 µm in each direction from the respective tonotopic frequency, 200 µm in total. For evaluation of ANF, the signal intensity of NF200kD was measured in a 100 µm wide segment at each frequency. The region of interest (ROI) was chosen from the level of IHC to the habenula perforata (**Fig. 4a**, dashed line). The size of the region of interest was the same in each image and brightness levels ranged from 0 to 4095 arbitrary intensity units (a.u.). Integrated density of each ROI was quantified using Image-J. For IHC synapses, two adjacent tiles of IHCs were imaged at 62x magnification in Z-Stacks in 0.5 µm steps. Intact and orphaned IHC synapses were visualized by co-labeling presynaptic CtBP2 positive ribbons and postsynaptic GluA2 terminals. Total intact synapses and orphaned ribbons per IHC were counted using ImageJ’s “Cell Counter” plugin. All specimen were evaluated in a blinded fashion to the treatment and frequency by pseudo-anonymizing the labeling prior to counting.

### 5. RNA isolation and RT-qPCR in an organotypic explant model of EIT

mRNA expression of pro-inflammatory cytokines and enzymes were evaluated using an adapted version of a previously described *in vitro* cochlear explant EIT model [24]. Cochleae of 4-day old (P4) C57BL/6 mice were extracted following rapid decapitation and divided into three groups: (1) a cochleostomy-only group (Control), (2) an EIT group, mimicked by the insertion of a monofilament suture through a cochleostomy, and (3) EIT with an additional 10 µM of AC102 for 24 hours during incubation (**Fig. 1c)**. In both trauma groups, a 2-0 non-resorbable monofilament suture (Ethilon, Ethicon) was inserted 3 times through an enlarged RW to enable a larger insertion angle. Cochleae were then incubated in PBS for 10 minutes with the suture *in situ*. Subsequently, cochleae were dissected as Organ of Corti explants and split into a basal and middle-to-apical part. In each group with each experiment, a total of 6 apical or basal specimens were pooled and incubated in the same well for 24 hours using Dulbecco’s modified Eagle’s medium (DMEM; Invitrogen, Carlsbad, CA, USA) containing glucose (4.5 g/L), 1% of N-1 supplement (Sigma Aldrich), and 1% ampicillin (Sigma Aldrich). For the AC102 treatment, 10 µM of AC102 was additionally added to the culture medium.

Total RNA was extracted using the RNeasy Mini Kit (Qiagen, Hilden, Germany) according to the manufacturer’s protocol. Tissue was transferred into a sterile 2 ml tube containing 600 µl of lysis buffer with added β-mercaptoethanol. Then, samples were homogenized using an ultrasonic homogenizer (Hielscher Ultrasonics, Teltow, Germany) to maximize the yield and quality of total RNA. Off-column DNase treatment of all inner ear samples was carried out using the RNase-Free DNase Set (Qiagen). Quantity and quality of the obtained total RNA samples were analyzed using a spectrophotometer (Nanodrop, Thermo Fisher Scientific, Waltham, MA, USA). 300 ng of total RNA was considered appropriate for cDNA synthesis using the RevertAid H Minus First Strand Synthesis Kit (Thermo Fisher Scientific). Reverse transcription was performed for 60 min at 42°C, 5 min at 95°C, and cooling to 4°C using a thermocycler (Avantor VWR, Radnor, PA, USA). All primers were originally designed using NCBI primer blast software. Primer sequences for forward and reverse primers used in this study are depicted in **Suppl. Table 1**. RT-qPCR was performed in translucent 96-well plates using the Applied Biosystems 7500-Real-Time PCR System (Applied Biosystems, Waltham, MA, USA). For each well, the 20 µl reaction contained: 10 µl of Luna® Universal qPCR Master Mix, 0.5 µl of forward and 0.5 µl of reverse primers, 5 µl of cDNA template and 4 µl of RNAse-free water. The cycling conditions were: preincubation at 95 °C for 1 min followed by 45 cycles of amplification including denaturation at 95°C for 15 seconds and annealing at 60 °C for 30 seconds. Upon PCR completion, melt curves at 60 °C to 95 °C were generated to check for contamination and primer specificity. No-template controls were carried out for each primer pair to check for presence of primer dimer formation. Additionally, “no-RT” controls lacking the reverse transcriptase were performed to rule out the presence of genomic DNA contamination.

The amplification efficiency for each primer pair was estimated using serial dilutions of the obtained cDNA samples with all samples run in duplicates. Serial dilution series and RT-qPCR efficiency was calculated from the slope according to the equation: Efficiency = (10(-1/slope) – 1) x100. Criteria for an optimal qPCR reaction were: (1) efficiency within the range of 95%-105%, and (2) well-defined melting curves with a single product-specific melting temperature. Using the 2-ΔΔCq method [41], relative changes in mRNA gene expression were analyzed and normalized to the reference gene β-actin as well as to the obtained control samples, and presented as mean-fold units (mfu).

### 6. Ethanol and AC102 treatment of an HT22 cell line

Anti-apoptotic activity of AC102 treatment were evaluated in an ethanol-challenged immortalized mouse hippocampal cell line (HT22) by evaluation of cell morphology as well as caspase 3/7 activity *in vitro*. In short, a previously described apoptosis-inducing ethanol (EtOH) assay using 4.5% EtOH treatment for 5 hours was utilized (**Fig. 1d)** [42]. For morphological analysis, 40,000 HT22 cells per well were seeded in 24-well plates and maintained under CO_2_ at 37^0^C in DMEM medium containing 10% FCS for 24 hours. Afterwards, culture medium was replaced with fresh DMEM medium containing 4.5% EtOH alone or in combination with increasing concentrations of AC102 (1, 10, 30 or 100 μM), and incubated for another 5 hours. Following the incubation, 8-10 images were acquired from 2 wells per group by bright field microscopy (Zeiss Axiovert 10). Damaged cells, identified by a round shape with fragmented membranes (**Fig. 6c**), were counted and reported as percentage (%) of all cells. For analysis of caspase 3/7 activity, 5,000 – 10,000 HT22 cells per well were seeded in 96-well plates and maintained under similar conditions to the morphological experiments. After 24 hours of incubation in culture medium, cells were similarly treated with 4.5% EtOH, and co-cultured with 30 μM or 100 μM of AC102 for another 5 hours. Caspase activity was measured with the Caspase-Glo® 3/7 (Promega Corp., Fitchburg, WI, USA) assay kit, according to the manufacturer’s instructions. For immunocytochemistry, following the same treatment conditions with 4.5% EtOH and co-culture with additional AC102 (30 μM and 100 μM), cells were fixed with 4% PFA, permeabilized with 0.2% Tween-20 in PBS and blocked with 0.2% Tween-20 in PBS containing 5% NGS. Cells were then immunostained with Cleaved Caspase-3 antibody (CST9661, Cell Signaling Technology, Massachusetts, USA) at 1:200 dilution followed by the appropriate secondary Alexa-Fluor-488 antibody (A-11034, Invitrogen) at 1:500 dilution. Nuclei were stained with Hoechst dye. Images were acquired using a Zeiss fluorescence microscope at 40x magnification. The integrate densities were quantified using Image-J. For each of 7 biological replicates, 10-15 images were quantified per treatment group.

#### Statistics and calculations

Data in the text is presented as mean ± standard deviation (SD). Data points in graphs represent mean values. Error bars represent the SD or standard error of the mean (SEM) as indicated in the respective figure legend. Data analysis was performed using GraphPad Prism 9.5 (GraphPad Software, San Diego, CA, USA). Statistical analysis of electrophysiological data was performed using unpaired t-tests. Statistical differences in histological data were evaluated by unpaired t-tests or one-way ANOVAs with Tukey’s post-hoc analysis, as suitable for the respective dataset. For statistical analysis of RT-qPCR data and apoptosis data, significance between the individual groups was tested using one-way ANOVAs with Tukey’s post-hoc analysis, or repeated one-way ANOVA with Dunnett’s post-hoc, respectively. Results were considered statistically significant if the p-value was lower than 0.05.

## Results

### AC102 promotes earlier recovery of auditory thresholds following cochlear implantation

To determine the protective effects of AC102 on EIT, AC102 (n = 9) or a Vehicle hydrogel (n = 9) were applied into the RW niche of female Mongolian gerbils 24 hours prior to CI. Changes in auditory function were investigated by auditory CAPs over 28 days following CI (**Fig. 1a**).

Electrode insertion resulted in a significant increase of CAP thresholds across all measured frequencies (p<0.005) compared to their preoperative baseline measurements (**Suppl. Fig. 1**). Threshold shifts in the apical cochlear regions were smallest, and frequencies just apical of the maximum insertion depth, showed a noticeable increase (**Fig. 2a-b**). The most prominent changes were observed between 6.3 and 32 kHz, which tonotopically represents the area directly affected by the insertion of the electrode (**Fig. 2c-d**). Acute postoperative threshold shifts were similar between both groups. Across all frequencies, threshold shifts in the Vehicle group increased until day 7 before recovering, whereas AC102-treated animals showed an earlier recovery after postoperative day 3. In the frequency range from 0.5-1.5 kHz (**Fig. 2a**), significantly smaller threshold shifts were seen from day 14 in AC102-treated group (p=0.017). At 2.0-32 kHz (**Fig. 2b-d**), threshold shifts became significantly smaller on day 7 in the AC102 group. This significant difference continued across all cochlear frequencies until day 28. At 0.5-1.5 kHz and 2.0-5.0 kHz, a difference of 12.6 ± 6.1 to 22.7 ± 10.7 (p=0.016) and 37.9 ± 10.5 to 56.9 ± 8.5 dB (p=0.0002) was observed in the AC102-treated animals compared to the vehicle control group, respectively (**Fig. 2a-b**). Despite larger inter-individual variability in the high frequencies, similarly significant differences of 56.0 ± 11.4 dB to 66.1 ± 4.2 dB (p=0.012) and 57.5 ± 22.0 dB to 74.3 ± 6.7 dB at (p=0.012) were observed at 6.3-12.6 kHz and 16-32 kHz, respectively (**Fig. 2c-d**).

Electrode impedances increased in both treatment groups from day 7 until day 28, primarily at the basal electrode contact. AC102 treatment showed a tendency towards lower impedances on the apical contact compared to the Vehicle treatment, without revealing statistically significant differences (**Suppl. Fig. 2**).

### AC102 significantly preserves HCs and neuronal connections, and narrows cochlear trauma to directly mechanically affected regions

To confirm the beneficial effect of AC102 on a cellular level, cochleae were histologically processed and analyzed. Implanted cochleae (AC102, n = 9; Vehicle, n = 7) and randomly selected non-implanted, contralateral cochleae of the Vehicle group (Control, n = 4), representing an untreated control, were prepared as cochlear whole mounts, and analyzed after immunofluorescence staining (**Fig. 3a**).

**Fig. 3.**
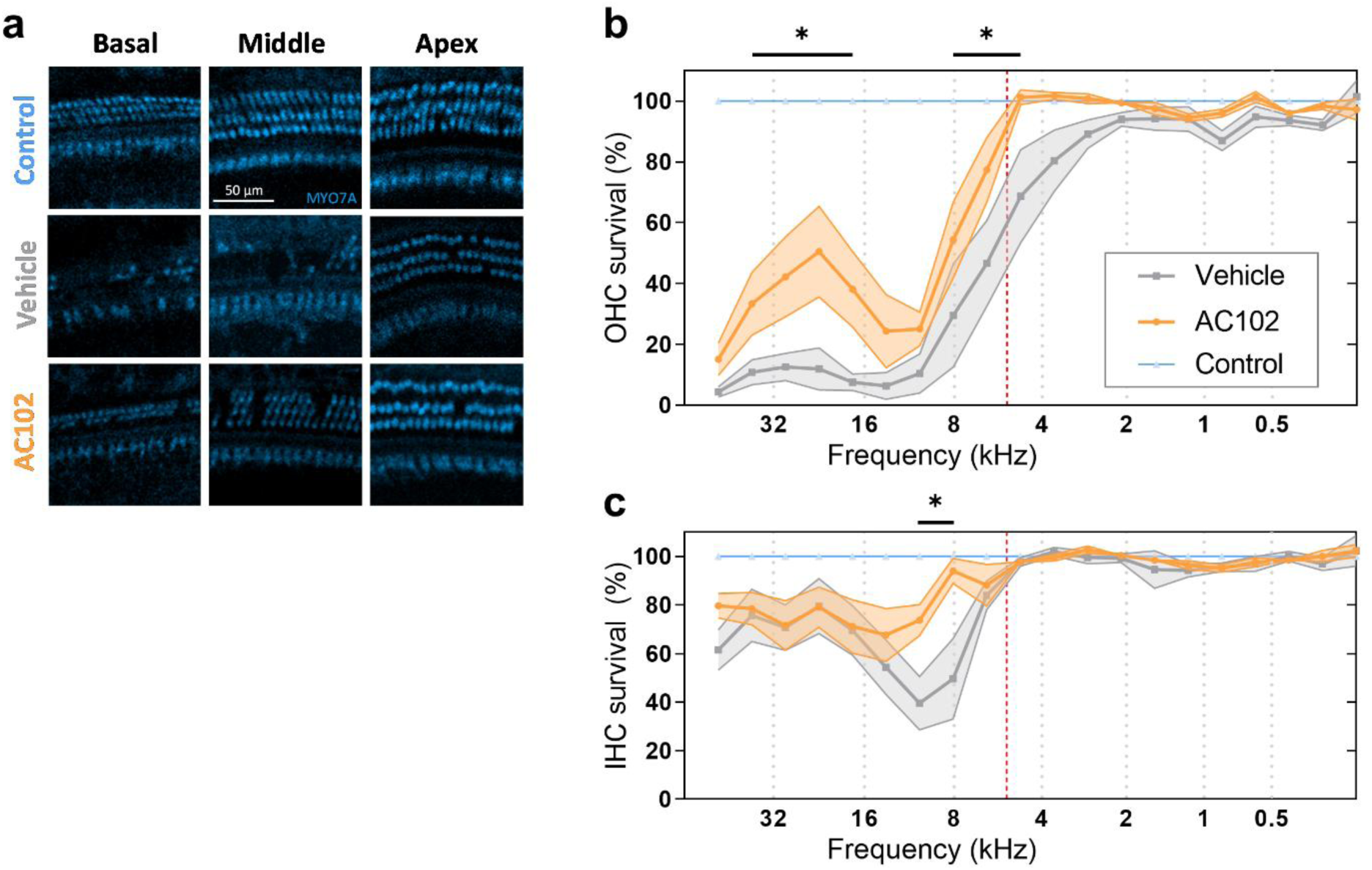
Quantification of outer and inner hair cell loss in cochlear implanted Mongolian gerbils treated with AC102 or Vehicle control hydrogel. The effect of a preoperative administration of AC102 versus Vehicle control 24 hours prior to surgery was investigated on a histological level. After the last follow-up measurement 28 days after cochlear implantation, cochleae were extracted for immunofluorescence analysis to determine the degree of OHC and IHC loss. **a**. Illustrative images from the basal (left column), middle (middle column), and apical (right column) cochlear areas (scale bar, 50 µm), stained with myosin VIIa (MYO7A, blue) for hair cell cytoplasm of the Control (1^st^ row), Vehicle (2^nd^ row) and AC102-treated (3^rd^ row) groups. Cytocochleogram of OHC (**b**) and IHC (**c**) survival in the AC102 (n = 9, orange) and Vehicle-treated (n = 7, gray) group in % compared to the non-implanted contralateral sides (blue), shown as mean and standard error of the mean. A frequency-map of the gerbil was used to map the hair-cell segments to specific frequencies. The red-dashed line indicates the maximum insertion depth of the electrode. * = p < 0.05

CI resulted in a pronounced loss of OHCs in the basal part of the cochlea, the region of the inserted electrode (**Fig. 3b**). Vehicle-treated cochleae had a near-complete loss of OHCs (only 4-13% preserved cells) while AC102-treated animals showed significantly higher remaining OHCs (33-50%, p=0.04 – 0.0006). A gradual increase of OHCs was observed in the Vehicle group from 30% at 8.0 kHz to 93% at 2.0 kHz. The number of OHCs in the AC102-treated cochleae was significantly higher with 54% remaining OHCs from 8.0 kHz up to 100% at 6.0 kHz (p=0.026 – 0.0037). Apical to the maximum insertion depth (**Fig. 3b**, red dashed line), OHCs were fully preserved in AC102-treated cochleae compared to the Vehicle group. IHCs were more robust and displayed a maximum loss in the area just basal of the maximum insertion depth (**Fig. 3c**). From 12 to 8.0 kHz, loss of IHCs was significantly higher in the Vehicle group (39% and 50%) compared to the AC102-treated group (74% and 94%, p=0.013). In the most basal region, 61-80% of IHCs irrespective of the group were preserved. Apical to the maximum insertion depth, IHCs were completely preserved in both groups.

Further, neuronal, and synaptic loss was quantified in one-octave steps from 32 to 0.5 kHz (**Fig. 4a**). Loss of type-II ANFs was most pronounced between 32 and 8.0 kHz in Vehicle-treated animals. AC102-treated cochleae showed significantly higher numbers of type-II ANFs per 200 µm at 32 kHz (22.0 ± 7.6 to 13.0 ± 4.5, p=0.002), and 16 kHz (19.2 ± 7.6 to 10.9 ± 7.0, p=0.004), compared to the Vehicle group. From 8.0 to 1.0 kHz, higher numbers of type-II ANFs in the AC102-treated cochleae compared to the Vehicle group were noticed, without reaching statistical significance. At 0.5 kHz, type-II ANFs were significantly lower in the Vehicle-treated group compared to the AC102-treated cochleae (11.3 ± 4.3 to 23.1 ± 3.7, p<0.001, **Fig. 4b**).

**Fig. 4.**
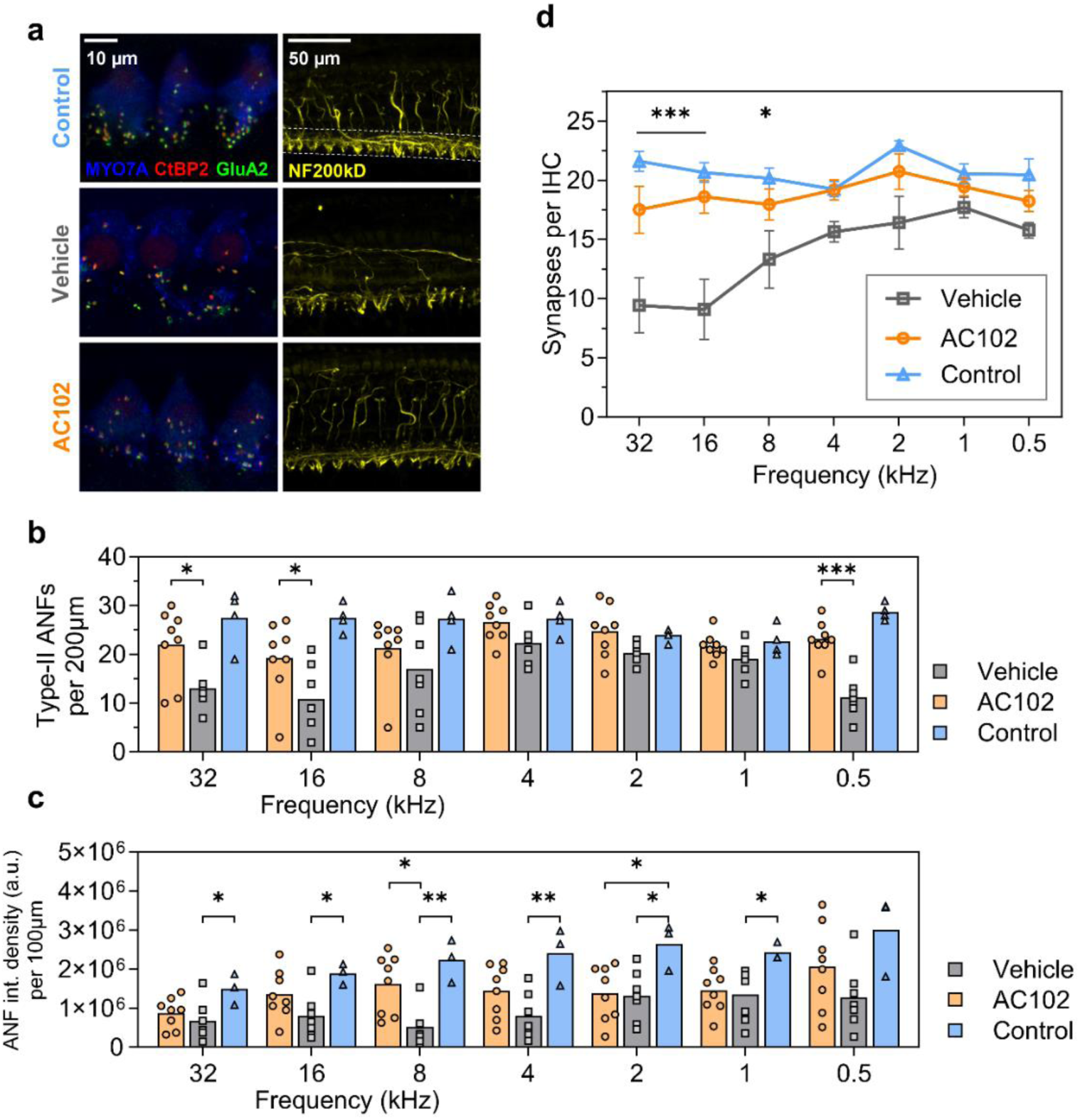
Quantification of neuronal elements of cochlear implanted gerbils treated with AC102 and Vehicle control. **a.** Exemplary images of inner hair cell synapses (left row, scale bar, 10 μm) and ANFs (right row, scale bar, 50 µm) from the 16 kHz region of the Vehicle and AC102-treated animals and untreated control group. The presynaptic ribbon was stained with CtBP2 (red), which also faintly labels IHC nuclei. The postsynaptic synapse was visualized with GluA2 (green). Hair cell bodies were visualized by staining of Myo7A (blue) and auditory nerve fibers by NF200kD (yellow). Dashed lines indicate the area for measurement of integrated density in a range of 100 µm. **b**. Absolute survival of type-II ANFs (numbers per 200 µm) in the frequency ranges from 32 and 0.5 kHz of the AC102 (n = 8, orange), Vehicle-treated (n = 7, gray), and contralateral untreated (n = 4, blue) groups. **c.** Signal intensity (integrated density) of ANF in a 100 µm wide area from the inner hair cell to the habenula perforata. **d**. Absolute number of intact synapses per IHC between 32 kHz and 0.5 kHz in one-octave steps on the endpoint day 28 of the AC102 (n = 6, orange), Vehicle-treated (n = 6, gray), and contralateral untreated (n = 4, blue) groups. * = p < 0.05, ** = p < 0.01 *** = p < 0.001 between AC102 and Vehicle group.

ANF signal intensity (**Fig. 4a**, dashed line) was significantly reduced in the Vehicle-treated group from 32 kHz to 1.0 kHz compared to the non-implanted control group (p<0.05). This difference was most pronounced at 8.0 kHz (5.09x10^5^ to 2.23x10^6^, p=0.004) and 4.0 kHz (7.94x10^6^ to 2.4x10^6^, p=0.007). AC102-treated animals showed a significantly higher signal intensity at 8.0 kHz compared to the Vehicle-treated group (1.62x10^6^ to 5.09x10^5^, p=0.011, **Fig. 4c**) and a significant reduction at 2.0 kHz (1.38x10^6^ to 2.65x10^6^, p=0.033) compared to the non-implanted control.

IHC synapses ranged from 21.6 ± 1.7 synapses per IHC at 32 kHz to 20.5 ± 2.6 at 0.5 kHz in the Control group. CI resulted in a pronounced loss of IHC synapses in the Vehicle-treated group at 32 kHz (9.4 ± 5.7) and 16 kHz (9.1 ± 6.2) and a partial loss between 4.0 to 0.5 kHz (15.6 ± 2.1 to 15.8 ± 1.4). Treatment with AC102 showed a near complete preservation of IHC synapses across the whole cochlear length, with significantly higher numbers at 32 kHz (17.5 ± 4.9, p<0.001), 16 kHz (18.6 ± 3.4, p<0.001) and 8 kHz (17.9 ± 3.2, p=0.049), compared to the Vehicle-treated group (**Fig. 4d**).

### AC102 decreases the expression of pro-inflammatory cytokines in an *ex vivo* model of electrode insertion trauma and reduces apoptosis in an ethanol challenged HT22 cell line

To evaluate the anti-inflammatory effects of AC102, mRNA expression of TNF-α as well as the inflammatory enzymes (iNOS and COX-2) were quantified in a modified version of a previously described *ex vivo* EIT model [24] (**Fig. 1b**). Overall, EIT resulted in an upregulated mRNA expression of TNF-α and iNOS, with a significant increase of iNOS in the apical part (p=0.025) compared to an untreated control group. COX-2 expression was unchanged following EIT. In the middle-to-apical region (**Fig. 5a**), treatment with 10 µM AC102 resulted in a significant lower expression of TNF-α (0.63 ± 0.2 mfu to 1.14 ± 0.3 mfu, p=0.04) and iNOS (1.04 ± 0.9 mfu to 4.95 ± 2.9 mfu, p=0.018) compared to the EIT only group, and a non-significant lower expression of COX-2 (0.69 ± 0.7 mfu to 0.97 ± 0.5 mfu). Similarly, AC102-treatment resulted in a significantly reduced expression of TNF-α (0.55 ± 0.2 mfu to 1.12 ± 0.5 mfu, p=0.02) and iNOS (0.55 ± 0.5 mfu to 1.64 ± 0.7 mfu, p=0.04) (**Fig. 5b**) in the basal part, compared to the EIT only group, whereas COX-2 showed a non-significant upregulation (1.82 ± 1.0 mfu to 1.05 ± 0.54).

**Fig. 5.**
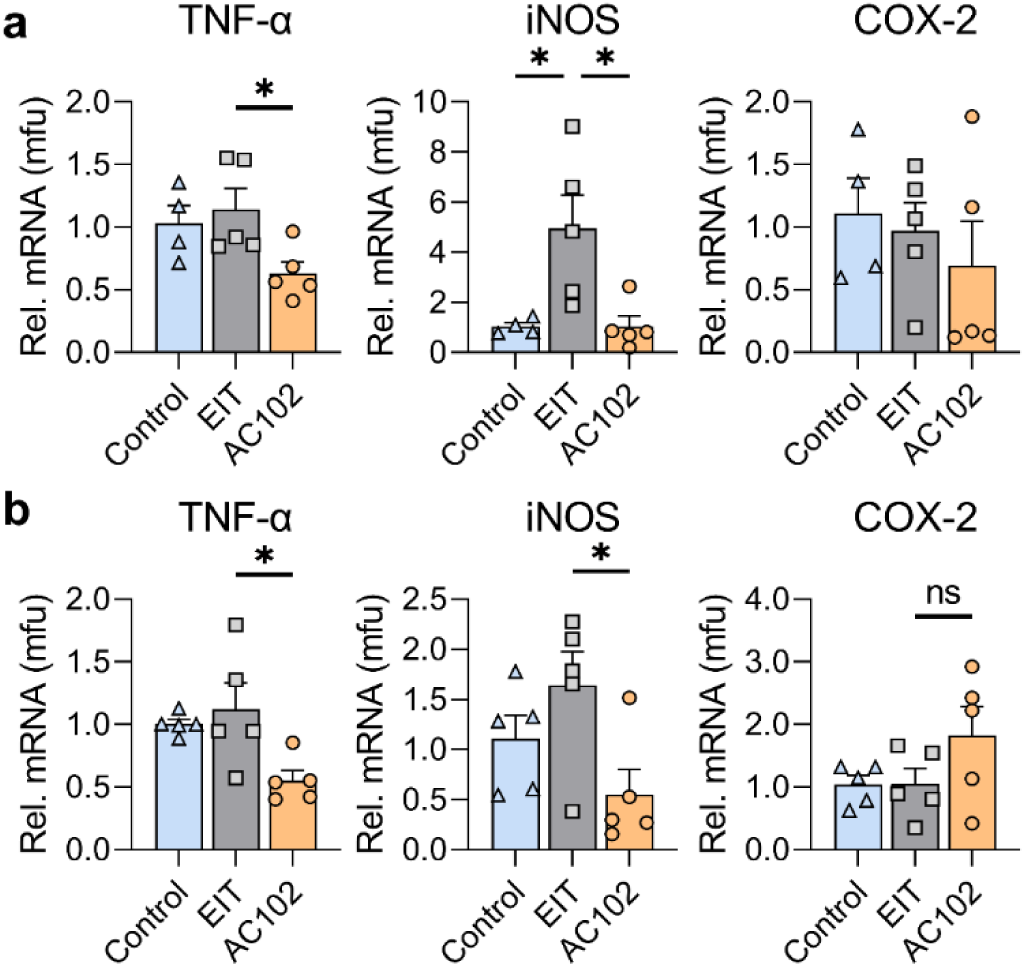
mRNA expression changes of pro-inflammatory cytokines 24 hours after electrode insertion trauma *ex vivo*. Electrode insertion in cochlear implantation was mimicked by inserting a 2-0 monofilament suture into an enlarged round window of P4 C57BL/6 mice cochleae (EIT, gray bar), and compared to a non-implanted group (Control, blue bar) and a 10 μM AC102-treated group after EIT (AC102, orange bar). **a.** The first row corresponds to the basal cochlear segment of each gene of interest. **b.** The second row corresponds to the same genes in the middle-to-apical cochlear segments. Error bars = SEM, n = 4-5 / group, * = p < 0.05, ns = no statistical significance

Further, the anti-apoptotic properties of AC102 were elucidated in an established ethanol-induced apoptosis assay *in vitro* (**Fig. 1c**). Ethanol challenging of HT22 cells significantly increased cell damage compared to untreated control (82.4 ± 11.48% to 9.0 ± 1.0%, p<0.0001), and resulted in an increase of caspase 3/7 activity (141.6 ± 75.94 a.u. to 66.76 ± 46.46 a.u., p<0.0001, **Fig. 6a,b**). Treatment with AC102 showed a concentration-dependent decrease of morphological cell damage, with a significant decrease at 30 µM (66.5 ± 9.51%, p=0.028) and highly significant decrease at 100 µM (35.99 ± 9.51; p<0.0001; **Fig. 6a**). A similar concentration-dependent reduction of caspase 3/7 activity was observed after treatment with AC102, significantly at 100 µM of AC102 (83.9 ± 63.23 a.u., p=0.0018, **Fig. 6b**). On a protein level, the expression of Cleaved-Caspase-3 similarly increased following EtOH-treatment, compared to the untreated control (6.60x10^6^ a.u. to 2.79x10^6^ a.u., p=0.0003). Co-culture of 100 µM AC102 significantly decreased Cleaved-Caspase-3 expression (3.88x10^6^ a.u, p=0.0065, **Suppl. Fig. 3**), similar to morphological and biochemical results.

**Fig. 6.**
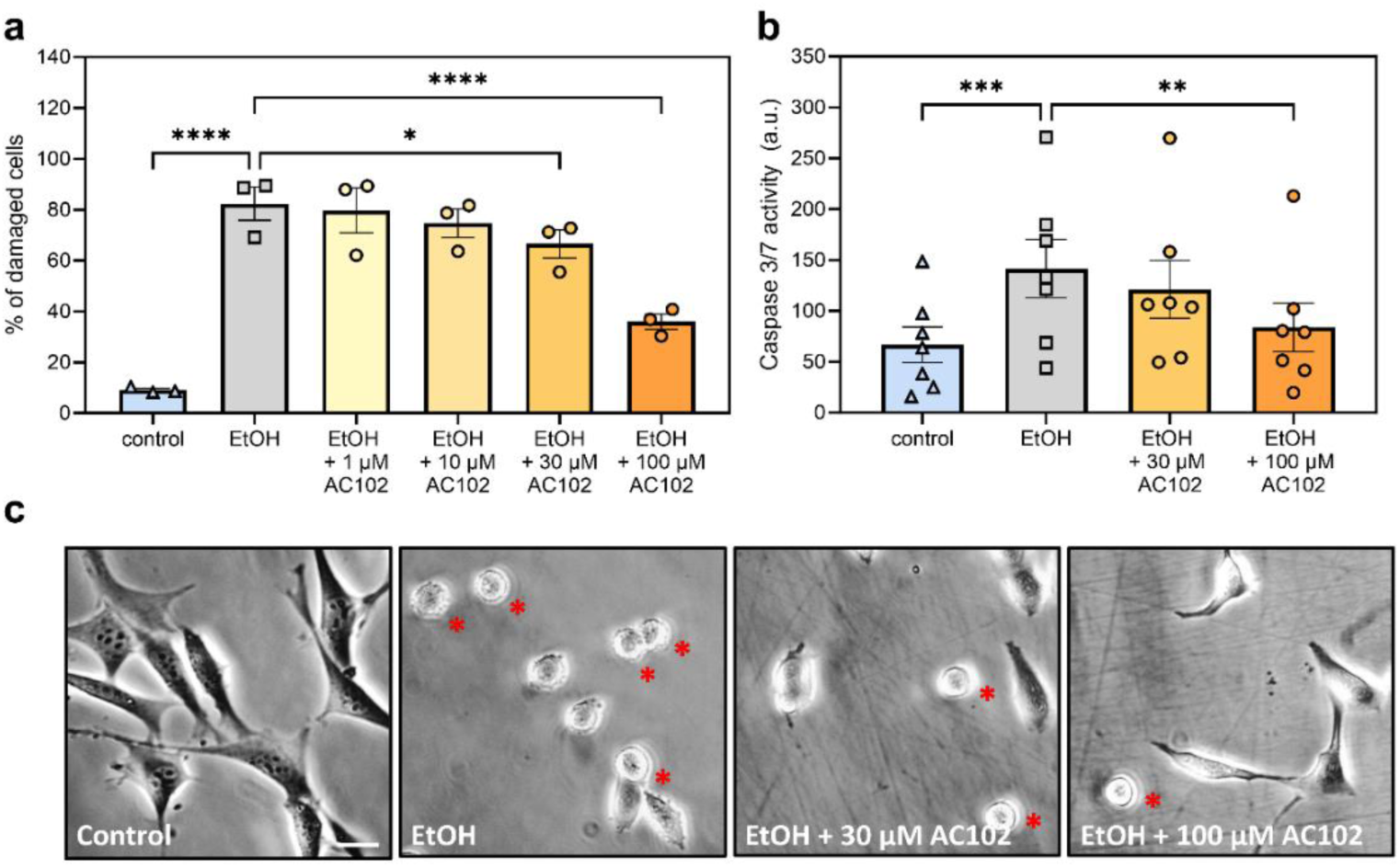
Evaluation of the anti-apoptotic effects of AC102 in ethanol challenged HT22 cells *in vitro*. HT22 cells were treated for 5 hours in cell culture medium (Control), added 4.5% ethanol (EtOH), and co-treatment with different concentrations of AC102. **a.** Percentage (%) of round cells following EtOH challenging with co-treatment of AC102 at 1, 10, 30 and 100μM (n = 3). **b.** Confirmation of AC102s anti-apoptotic effects by quantification of Caspase-3/7 activity using the Caspase-Glo® 3/7 assay. Arbitrary relative luminescence units (a.u.) were measured after a 5-hour incubation in culture medium, 4.5% EtOH and added 30 or 100 μM of AC102 (n = 7). **c.** Exemplary images of HT22 cells following the incubation with culture medium, added 4.5% EtOH and the additional 30 and 100 μM of AC102 (red asterisks indicate round damaged cells, scale bar = 20 µm). * = p < 0.05, ** = p < 0.01, *** = p < 0.001, **** = p < 0.0001

## Discussion

While the number of people with hearing impairment is projected to increase significantly in the coming years [1], only one pharmacological treatment option for a single etiology of inner ear trauma is currently approved [3]. In EAS-CI surgery, GCs are still routinely used due to their potent anti-inflammatory and immunosuppressive effects. However, recent meta-analyses and randomized control trials in humans failed to show a significant effect on hearing preservation [33], highlighting an unmet need for novel treatment options to counteract the progressive loss of residual hearing. Here, we demonstrate the otoprotective effects of AC102, a novel pyridoindole, in an animal model of CI, and elucidate on its mechanisms in *in vitro* and *ex vivo* experiments.

Electrode insertion resulted in immediate auditory threshold shifts across all frequencies in a basal to apical gradient, consistent to previous EITs in a variety of rodent species [30,32,43,44]. While in human patients’ surgeons follow principles to minimize EIT and the subsequent immune response, we intentionally opted for a traumatic insertion to decrease the possibility of masking the protective effects of AC102 due to spontaneous recovery of thresholds [45,46].

Immediately after CI, the 24-hour preoperatively applied AC102 had only a minor and non-significant effect on the preservation of auditory function compared to the control group. However, we observed a continuous recovery of auditory threshold shifts in the AC102-treated group over the subsequent course of the experiments, along the whole frequency range. In the Vehicle group, thresholds deteriorated for 7 days following CI and subsequently recovered slightly until day 28. This temporal pattern, in agreement with previous studies [32,47], likely stems from an inflammatory response initiated by EIT lasting for a week before gradually ceasing, followed by a prolonged wound healing response and restoration of cochlear homeostasis [23,25]. TNF-α and IL-1ß, cytokines expressed early during inflammation, promote the migration of monocytes and upregulation of apoptotic pathways for up to 14 days, resulting in a continuous inflammatory environment after acute injury [24,48]. In the AC102-treated group however, auditory thresholds started to recover earlier from day 3 and showed significant improvement throughout until day 28. This suggests that AC102 does not exert immediate effects but rather acts as a modulator of the cochlear immune response following EIT.

Immunohistochemical evaluation of the cytocochleogram enabled us to describe the effects of CI on HCs in unprecedented detail. On a morphological level, a significant loss of OHCs was observed in the vicinity of the electrode in the basal part of the cochlea. IHCs were more robust, yet reduced near the maximal insertion depth, likely due to the narrowing of the cochlear diameter towards the apex, which causes the inserted electrode to exert greater shear forces to the surrounding cochlear structures [49]. Previous studies of CI similarly observed a correlation between high auditory threshold shifts and loss of OHCs in the basal area [43,50]. Loss of residual hearing in the cochlear regions around the implanted electrode therefore likely resulted from the deterioration of HCs, which were significantly preserved in the AC102-treated group. However, animal models of CI with less severe insertion trauma did not indicate a correlation between threshold shifts and HC loss [30,45,51,52].

Interestingly, in the control group, the area of OHC loss extended apically beyond the maximum insertion depth. In this zone, which in EAS recipients represents the frequency range to be preserved, HCs might undergo secondary deterioration due to a propagating inflammatory response, without being mechanically affected by the implanted electrode. Programmed cell death initiated through different apoptosis pathways has been found to spread apically in the first 12 hours after CI and stays upregulated for at least 4 days [24,53]. Similarly, intense impulse noise, a sound-evoked trauma comparable to mechanical insertion, induces a localized activation of apoptosis in OHCs within 5 minutes, expanding from the initial lesion along the cochlea for 30 minutes after the event, supporting the theory of local spread of inflammation after insertion trauma [54]. AC102 treatment seemingly narrowed the cochlear trauma to the directly affected regions during implantation, with a near-complete survival of HCs just beyond the maximum insertion depth.

Despite observing low-frequency CAP threshold shifts in both groups, HCs were nearly completely preserved from 2 kHz to the cochlear apex, suggesting a different reasons for deterioration of hearing function. We therefore examined the cochlea’s neuronal structures to identify potential causes, given that research in age-related hearing loss and noise trauma suggest the ANFs and synapse between cochlear nerve and IHC to primarily degenerate following cochlear injury [55,56]. We observed a significant decrease of ANF signal intensity across the whole cochlea and a loss of type-II ANFs in the basal cochlear area, where loss of OHCs is most pronounced. While loss of neuronal structures in the basal region can be attributed to a secondary deterioration after the lost OHCs [57], apical ANFs might deteriorate primarily despite preservation of their inner hair cells. Similar findings have been observed in previous animal studies of cochlear implantation, arguing this reduction to aid to the loss of residual hearing in regions with complete preservation of HCs [32,45].

Our findings further revealed a 50-60% reduction of IHC synapses in the basal cochlear region and 20-25% across the remaining cochlea up to 500 Hz after CI. AC102 treatment resulted in a near-complete preservation of IHC synapses as well as a structural preservation of type-II ANFs across the whole cochlea. Following EIT, apoptosis has been observed in both HCs and supporting cells along the whole cochlea after 24 hours following injury [53]. Supporting cells ensure neuronal maintenance of IHC synapses through the continuous release of neurotrophins [58]. Additionally, the main mechanism of cochlear synaptopathy, glutamate excitotoxicity of postsynaptic terminals, is potentiated by the expression of TNF-α and IL-1ß [56,59,60]. Apoptosis of cochlear structures coupled with potentiating effects of pro-inflammatory cytokines might therefore result in a dominant loss of IHC synapses that appears to be prevented by AC102. Furthermore, AC102 showed neuro-regenerative properties in Rotenone challenged HT22 cells and IHC synapses following noise-induced hearing loss, highlighting the possibility of secondary regeneration, rather than primary protection [6]. Whether synapses were protected from immediate trauma or recovered over the following 28 days, however, cannot be distinguished in this experiment.

In summary, AC102s effects are likely elicited during the inflammatory phase of EIT and mediated by anti-apoptotic and anti-inflammatory properties. To confirm those likely modes of action, three *in vitro* assays were utilized. In our study, dose responsive anti-apoptotic effects of AC102 were observed in ethanol challenged HT22 cells on a morphological level, by measuring caspase 3/7 activity, as well as by evaluation of the protein-expression of cleaved caspase-3. EIT induces apoptosis via caspase-dependent pathways as well as mitochondrial stress [23], with similar characteristics to ethanol-induced apoptosis [42]. A concentration of 30 µM of AC102 was sufficient to significantly attenuate apoptosis. In line, Rommelspacher et al. observed a dose dependent decrease of ROS buildup in AC102-treated HT22 cells following Antimycin-A induction [6], attenuating an important apoptosis-pathway. Moreover, we utilized a previously reported organotypic *ex vivo* model of EIT [24] and investigated AC102’s effects on the expression of pro-inflammatory cytokines and enzymes following cochlear trauma. In contrast to previous experiments, our assay induced only a minor upregulation of TNF-α and pro-inflammatory enzymes iNOS and COX-2 following EIT [24,61]. Bas et al. [24] used albino rat pups, while we used C57BL/6 mice. Some studies suggest that unpigmented subjects are more susceptible to cochlear trauma such as ototoxicity, noise trauma, and presbycusis [62,63]. Melanosomes, significantly higher expressed in pigmented animals, are hypothesized to assist in buffering ROS [62]. The presence of more melanosomes in C57BL/6 mice could increase its resilience to EIT *ex vivo*, leading to a reduced immune reaction compared to previous studies [64]. Nevertheless, treatment with 10µM of AC102 could already significantly reduce mRNA expression of TNF-α and iNOS in apical and basal cochlear sections. Similarly, Hamann et al. observed a dose-dependent reduction of LDH release and caspase-3 activity in mesencephalic cell cultures treated with 9MP, underlining the anti-apoptotic and anti-inflammatory effects of pyridoindoles [7].

Taken together, a single preoperative application of AC102 significantly aided in the recovery of auditory function following CI and significantly protected against the loss of HCs apical to the implanted electrode, which is the most important zone to preserve in EAS-CI recipients. In addition, AC102 significantly preserved auditory nerve fibers and IHC synapses throughout the whole cochlea. Moreover, the inflammatory response in CI shares similarities with other etiologies of SNHL. Therefore, the protective effect in EIT may suggest a causal effect of AC102 and recommend it as a candidate for other causes of SNHL. Since AC102 is currently being studied in a Phase-II human trial in idiopathic SNHL, rapid translation for AC102 in EAS-CI recipients appears to be realistic [5]. However, further studies are needed to first examine the long-term effects of AC102 on hearing preservation in cochlear-implant recipients.

## Acknowledgements

The authors wish to thank Sandra Peiritsch for the keeping and taking excellent care of the animals. The authors also wish to thank AudioCure Pharma GmbH, for the timely provision of the AC102 and Vehicle hydrogels. The authors also thank Roland Hessler from MED-EL, Austria, for manufacturing and providing the gerbil electrodes.

## Conflict of Interest

**CA** is currently holding a grant from the Christian Doppler Research Association (CD Laboratory for Inner Ear Research) and is receiving funding from MED-EL Corporation, Innsbruck, Austria. **MN**, **EY**, **AJG** and **MG** were financed by this funding while conducting the studies. **LDL** receives research funding from Decibel Therapeutics and Amgen and has worked as an independent consultant for Gerson Lehrman Group and Conclave Capital. **SBr** and **PM** are employed by MED-EL as Senior Research Specialists and were involved in conceptualization of the study design. **RS** and **HR** are CEO and CSO of Audiocure Pharma GmbH, developing the compound AC102. Both had no involvement during conduction of the study or during analysis of the results. **MK** and **SB** are employees of Audiocure Pharma GmbH. The remaining authors declare no conflict of interest. The funding sources did not influence the study design, the interpretation of the data or the decision to publish this article.

## Author Contributions

**MN, CH, SBr, PM,** and **CA** contributed to conceptualization and writing of the manuscript; **MN, EY, MG, AJG** and **AMK** conducted the animal experiments, histological evaluation and the *ex vivo* EIT model; **SBe** conducted the *in vitro* apoptosis experiments; **MN, LDL, CH** and **CA** performed data and statistical analysis; **CH, MK, HR, RS, PM, LDL** and **CA** contributed to the supervision and interpretation of data. All authors contributed to the revision and editing of the manuscript. All authors approved the final version of the manuscript.

## Ethics Approval

Animal care, treatments and procedures were performed according to the guidelines of the Austrian Federal Ministry for Science, Research and Economy. All animal experiments were approved by the local animal welfare committee (66.009/0094-V/3b/2019).

## Funding

Conducted experiments were funded by a grant received from MED-EL Corporation, Innsbruck, Austria

## Data Availability

The data supporting the findings of this study are available from the corresponding author CA upon reasonable request. Restrictions apply to details of analytics and formulation of AC102.

## Notes

### Competing Interest Statement

Christoph Arnoldner is currently holding a grant from the Christian Doppler Research Association (CD Laboratory for Inner Ear Research) and is receiving funding from MED-EL Corporation, Innsbruck, Austria. Michael Nieratschker, Erdem Yildiz, Anselm J. Gadenstaetter and Matthias Gerlitz were financed by this funding while conducting the studies. Susanne Braun and Pavel Mistrik are employed by MED-EL as Senior Research Specialists and were involved in conceptualization of the study design. Reimar Schlingensiepen and Hans Rommelspacher are CEO and CSO of AudioCure Pharma GmbH. Both had no involvement during conduction of the study or during analysis of the results. Monika Kwiatkowska and Sujoy Bera are employees of Audiocure Pharma GmbH. Lukas D. Landegger receives research funding from Decibel Therapeutics and Amgen and has worked as an independent consultant for Gerson Lehrman Group and Conclave Capital.

