## Supplementary Information for "A novel pyridoindole improves the recovery of residual hearing following cochlear implantation after a single preoperative application"

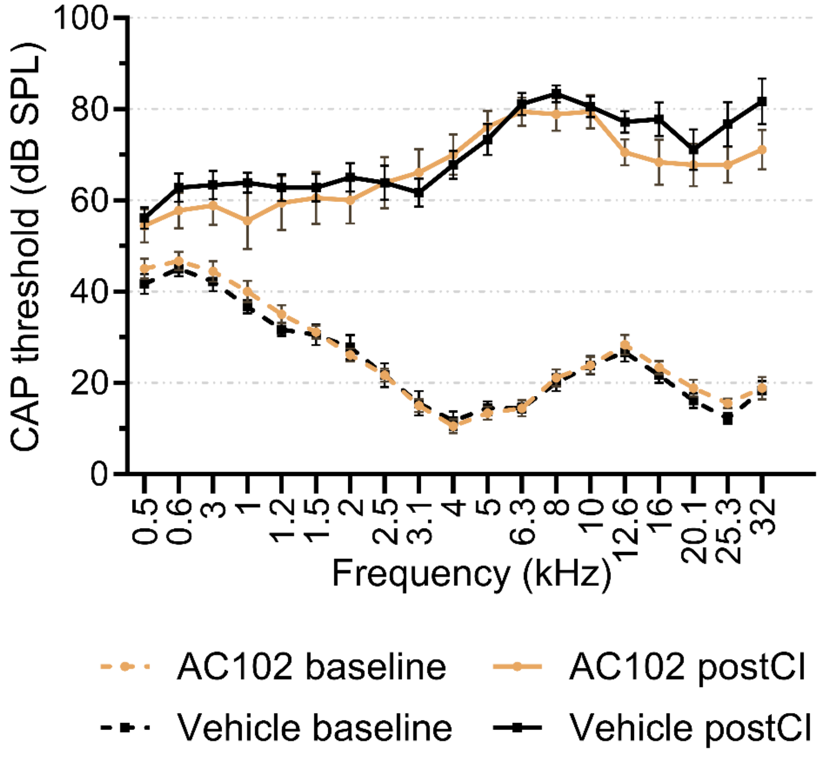


**Suppl. Fig. 1 Auditory thresholds (dB SPL) of AC102 (n = 9, orange) and Vehicle (n = 9, black) at the preoperative baseline measurement (dashed lines) and immediately after cochlear implantation (continuous lines).** A significantly elevated threshold was noted in each group at every measured frequency (p = 0.02 - < 0.0001); Error bars = SEM

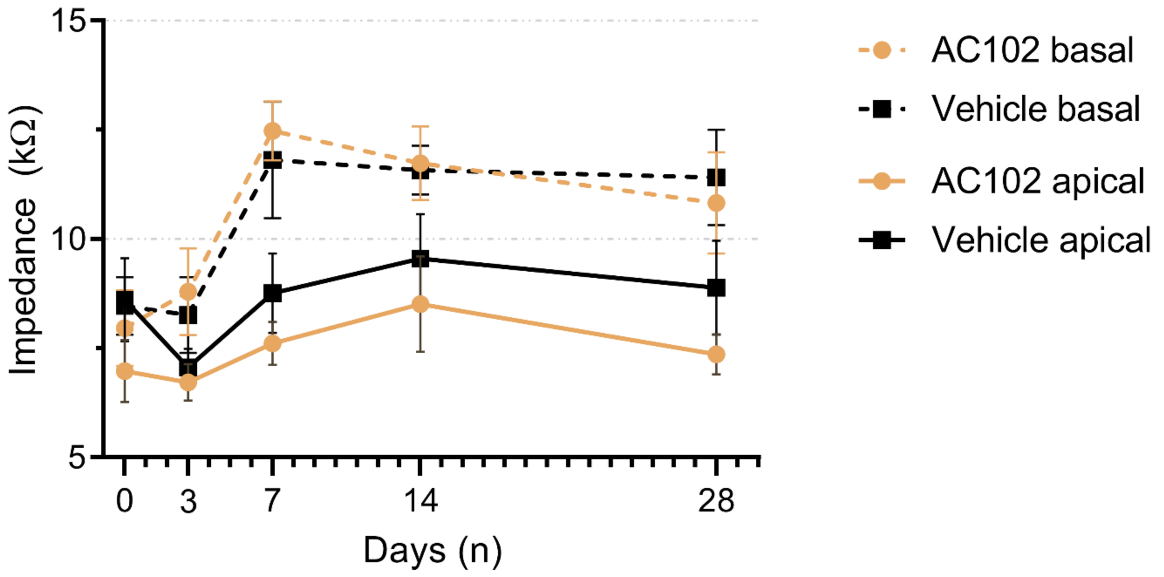


**Suppl. Fig. 2 Impedance changes of the basal (dashed lines) and apical (continuous lines) CI contact over the course of 28 days after cochlear implantation, treated with AC102 (orange) or Vehicle hydrogel (black) 24 hours prior to implantation.** Immediately following CI, impedance values were 6.97 ± 2.1 kΩ and 8.61 ± 2.8 kΩ on the apical electrode and 7.97 ± 2.6 kΩ and 8.47 ± 1.9 kΩ on the basal electrode in the AC102 and Vehicle animals, respectively, without any significant differences between both groups ($p > 0.05$; Error bars = SEM). For the apical contacts, impedances in the AC102 and Vehicle group increased from seven days after insertion to a maximum of 8.51 ± 3.2 kΩ and 9.56 ± 3.0 kΩ on day 14, before slightly decreasing to 7.36 ± 1.4 kΩ and 8.89 ± 3.2 kΩ on day 28. A trend towards lower impedances on the apical contact was observed in the AC102-treated group, whereas no significant difference was noted until day 28. At the basal electrode, impedances markedly increased 7 days after CI to 12.48 ± 2.0 kΩ and 11.81 ± 4.0 kΩ in the AC102 and Vehicle group and stabilized from there on. On day 28, impedances on the basal electrode were 10.83 ± 3.5 kΩ and 11.42 ± 3.3 kΩ in the AC102 and Vehicle group. No significant difference was observed between both groups ($p > 0.05$). Whereas other studies showed a progressive increase of impedance up to 90 days (34, 68), our study revealed a significant increase of impedances at the basal contact starting on day 7 and stagnated until day 28. This may suggest that acute insertion trauma, rather than a subsequent foreign body reaction, is the main trigger for an increase of impedances following CI in our study.

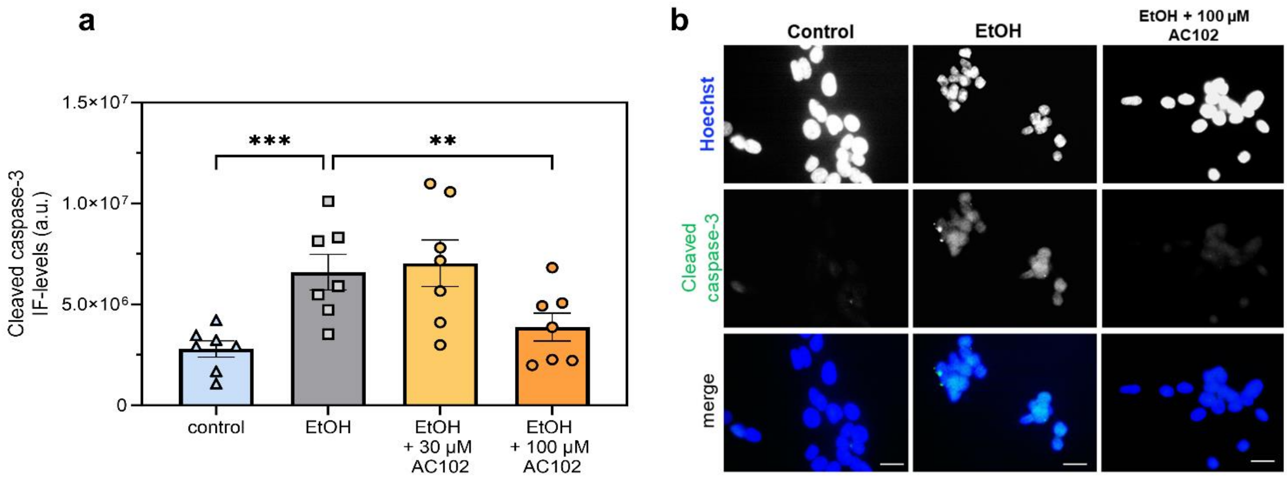


**Suppl. Fig. 3 Protein expression of cleaved caspase-3 in HT22 cells determined by immunofluorescence staining. a.** HT22 cells were treated for 5 hours in cell culture medium (Control), added 4.5% ethanol (EtOH), and EtOH with co-treatment of 30 µM and 100 µM of AC102 (n = 7). Cell nuclei were stained with Hoechst, and Integrated densities of cleaved caspase-3 expression measured as arbitrary units (a.u.). **b.** Exemplary panel of HT22 cells following the incubation with culture medium (left column), added 4.5% EtOH (middle column) and the additional 100 µM of AC102 (right column). Rows show example images of Hoechst (first row), cleaved caspase-3 (second row) and merged images (third row, scale bar = 50 µm). ** = $p < 0.01$, *** = $p < 0.001$

| **Target mRNA** | **Forward (5' to 3') primer** | **Reverse (3' to 5') primer** |
|---|---|---|
| β-Actin | CTCTGTGTGGATCGGTGGCT | CGCAGCTCAGTAACAGTCCG |
| TNF-α | CAGGCGGTGCCTATGTCTCA | GGCTACAGGCTTGTCACTCG |
| Cyclooxygenase 2 (COX-2) | ACAACATCCCCTTCCTGCGA | TGGGCAGTCATCTGCTACGG |
| Nitric oxide synthase 2 (NOS-2) | TCAGCCACCTTGGTGAAGGGA | GAAACTTCCAGGGGCAAGCCA |

**Suppl. Table 1 List of primer sequences used for evaluation of mRNA expression in the *ex vivo* model of electrode insertion trauma.**
